# Dysfunctional LHX6 pallido-subthalamic projections mediate epileptic events in a mouse model of Leigh Syndrome

**DOI:** 10.1101/2024.10.02.616229

**Authors:** Laura Sánchez-Benito, Joan Compte, Miquel Vila, Elisenda Sanz, Albert Quintana

**Affiliations:** Institut de Neurociències, Universitat Autònoma de Barcelona. Bellaterra (Barcelona) 08193. Spain; Department of Cell Biology, Physiology and Immunology, Universitat Autònoma de Barcelona. Bellaterra (Barcelona) 08193. Spain; Neurodegenerative Diseases Research Group, Vall d’Hebron Research Institute (VHIR)-Network Center for Biomedical Research in Neurodegenerative Diseases (CIBERNED); 08035 Barcelona, Spain; Department of Biochemistry and Molecular Biology, Autonomous University of Barcelona; 08193 Barcelona, Spain; Catalan Institution for Research and Advanced Studies (ICREA); 08010 Barcelona, Spain; Focus Area for Human Metabolomics, Faculty of Natural and Agricultural Sciences, North-West University, Potchefstroom, South Africa

**Keywords:** Epilepsy, Mitochondrial disease, Subthalamic nucleus, globus pallidus, *Lhx6*

## Abstract

Deficits in the mitochondrial energy-generating machinery cause mitochondrial disease (MD), a group of untreatable and usually fatal disorders. Among many severe symptoms, refractory epileptic events are a common neurological presentation of MD. However, the neuronal substrates and circuits for MD-induced epilepsy remain unclear. Here, using mouse models of mitochondrial epilepsy that lack mitochondrial complex I subunit NDUFS4 in a constitutive or conditional manner, we demonstrate that mitochondrial dysfunction leads to a reduction in the number of GABAergic neurons in the rostral external globus pallidus (GPe) and identify a specific affectation of pallidal *Lhx6*-expressing inhibitory neurons. Our findings further reveal that viral vector-mediated *Ndufs4* re-expression in the GPe effectively prevents seizures and improves the survival in the models. Additionally, we highlight the subthalamic nucleus (STN) as a critical structure in the neural circuit involved in mitochondrial epilepsy, as its inhibition effectively reduces epileptic events. Thus, we have identified a novel role for pallido-subthalamic projections in the development of epilepsy in the context of mitochondrial dysfunction. Our results suggest STN inhibition as a potential therapeutic intervention for refractory epilepsy in patients with MD providing new leads in the quest to identify novel and effective treatments.

## INTRODUCTION

Mitochondrial diseases (MD) constitute a group of genetic disorders characterized by alterations in mitochondrial function, primarily leading to defective oxidative phosphorylation (OXPHOS), affecting 1 in 4,300 births (Gorman et al. 2015). MD are often progressive, involving multiple organ systems. However, they primarily affect organs that rely on aerobic metabolism, such as the central nervous system (CNS), and muscular tissue. As a result, clinical manifestations are varied, but typically feature prominent neurological and muscular components (Gorman et al. 2016; McFarland, Taylor, and Turnbull 2010).

Among the different signs, epilepsy stands out as one of the most prevalent central nervous system manifestations in MD. Epileptic seizures lead to rapid neuronal damage, characterized by histopathological findings such as microvacuolation, neuronal loss, eosinophilia, astrogliosis, and secondary myelin loss in affected patients (Alston et al. 2017). Reports indicate that epilepsy occurs in 35-60% of individuals with biochemically-confirmed MD (Debray et al. 2007; Khurana et al. 2008). Early-onset epilepsy is associated with reduced life expectancy in childhood, with a 50% mortality rate within the year following the onset of epilepsy (Desguerre et al. 2014). Notable differences in epilepsy prevalence exist among various MD, and significant heterogeneity in the presentation of epileptic seizures among patients has been observed (Rahman 2012). These factors have contributed to our limited understanding of the pathogenic mechanisms underlying epilepsy in mitochondrial disorders.

Several hypotheses have been posited to link mitochondria dysfunction to epilepsy. For instance, energetic failure coupled with imbalances in calcium and ROS homeostasis could precipitate epileptic events, which in turn demand substantial energy. These events would subsequently exacerbate mitochondrial dysfunction, establishing a vicious cycle that ultimately leads to neuronal death (Rahman 2012; Desguerre et al. 2014; Kunz 2002). Alternatively, it has been proposed that the underlying pathophysiological process associated with mitochondrial epilepsies is initiated by the deficiency of the mitochondrial respiratory chain function in interneurons, which have been described to be highly susceptible to OXPHOS deficiencies, especially those resulting from defects in complex I and complex IV (Lax et al. 2016; Rahman 2018; Ticci et al. 2020). Thus, this would disrupt excitatory/inhibitory synaptic networks and promote neuronal hyperexcitability at the circuit level (Iizuka et al. 2002).

Several brain regions have been associated with epileptogenic circuitries, including the hippocampus, amygdala, frontal, temporal, and olfactory cortex, and basal ganglia (Vuong and Devergnas 2018; Chauhan et al. 2022). However, the discrete areas involved in mitochondrial epilepsy remain unclear. Therefore, a deeper understanding of the anatomical basis of epilepsy is crucial for comprehending the origin of abnormal electrical activity, facilitating accurate diagnosis, and guiding appropriate management strategies.

In this regard, basal ganglia lesions are a hallmark of MD (Eom et al. 2017; Schubert Baldo and Vilarinho 2020), and the basal ganglia are believed to play a direct role in seizure generation, with multiple studies demonstrating their significant involvement in the propagation and control of various seizure types (Vuong and Devergnas 2018; Deransart et al. 2001; Veliskova and Moshe 2006; Cheng et al. 2015). To explore the potential role of the basal ganglia in mitochondrial epilepsy, we dissected the epileptogenic circuit in an animal model of Leigh Syndrome, the most prevalent pediatric MD (Lake et al. 2016), which lacks mitochondrial complex I subunit NDUFS4 either ubiquitously or conditionally in GABAergic neurons (Kruse et al. 2008; Quintana et al. 2010; Bolea et al. 2019; Manning et al. 2023).

## RESULTS

### Restoration of *Ndufs4* in external pallidal GAD2 neurons extends lifespan, reduces epileptic events and abolishes local microglial reactivity in Ndufs4cKO mice

Mice lacking the NDUFS4 subunit of mitochondrial complex I in *Gad2*-expressing GABAergic neurons (Ndufs4cKO mice) exhibit epileptic seizures, reduced lifespan, and glial reactivity in the external globus pallidus (GPe) within the basal ganglia (Bolea et al. 2019; Manning et al. 2023). Thus, we aimed to investigate the potential involvement of mitochondrial dysfunction in the GPe in the development of epilepsy by using viral vector-mediated stereotaxic restoration of NDUFS4 in this area. The rescue of *Ndufs4* expression in the GPe of Ndufs4cKO mice (Ndufs4cKO-vr mice, rescue group) significantly increased lifespan compared to those injected with the control viral vector (Ndufs4cKO-vYFP, Figure 1A-C).

**Figure 1.**
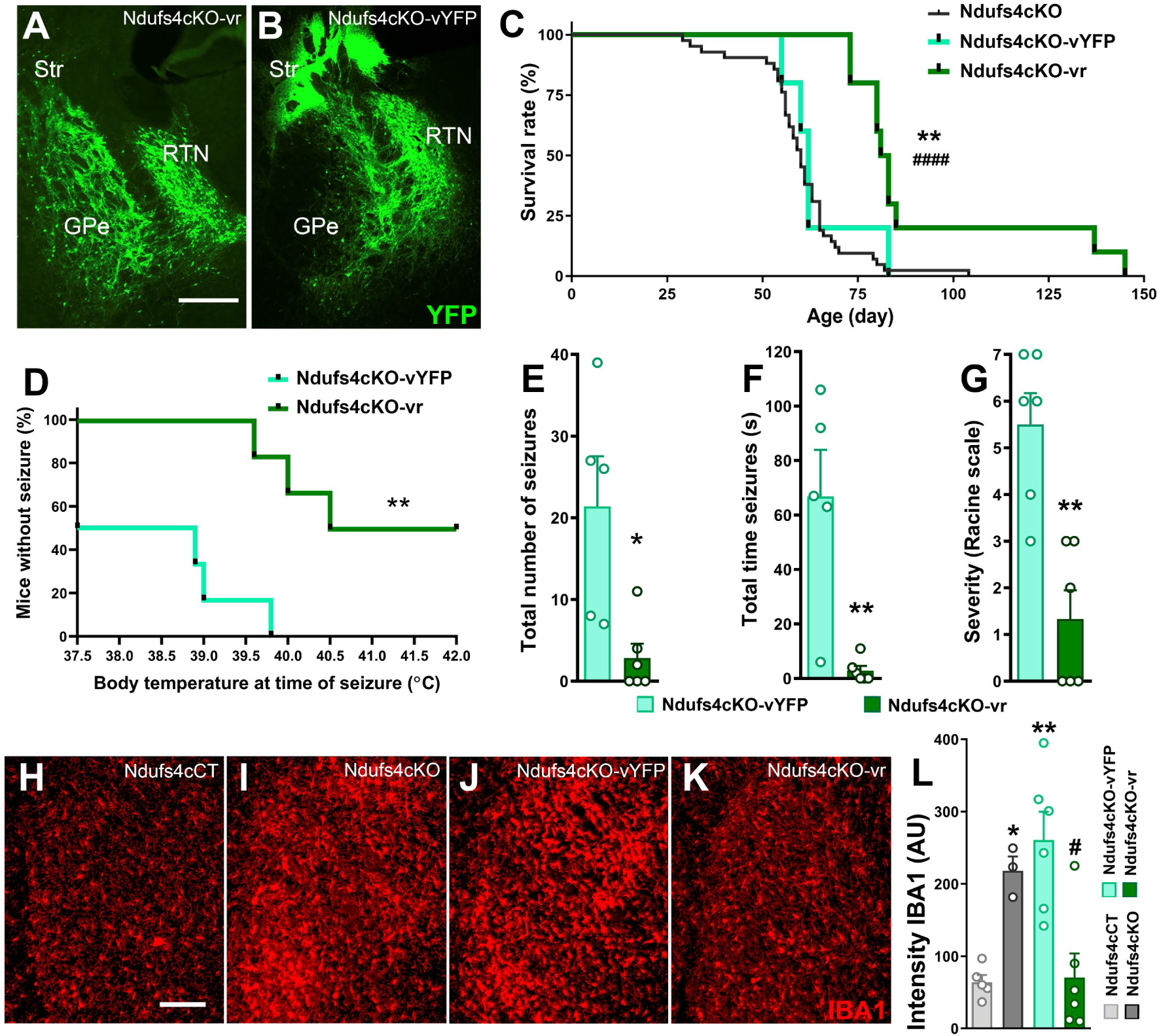
Re-expression of *Ndufs4* in *Gad2-*expressing GPe neurons reduces epilepsy and local microgliosis. **A-B)** Representative images showing the injection site in the GPe of Ndufs4cKO-vr (A) and Ndufs4cKO-vYFP (B) mice. Scale bar=400 µm. Str: Striatum; GPe: Globus Pallidus externus; RTN: Reticular Thalamic Nucleus. The reduced transduction observed in the GPe of Ndufs4cKO-vYFP is likely due to neuronal death resulting from NDUFS4 deficiency, as shown in Fig. 2. **C)** Survival curves for Ndufs4cKO-vYFP (control group, n=5), Ndufs4cKO-vr (rescue group, n=7), and uninjected Ndufs4cKO mice (n=42). ** p<0.01 indicates significant differences compared to Ndufs4cKO-vYFP group. ### p<0.001 indicates significant differences compared to the uninjected Ndufs4cKO group. Gehan-Breslow-Wilcoxon test. **D)** Percentage of animals without epilepsy relative to body temperature. n=6 per group. ** p<0.01, Gehan-Breslow-Wilcoxon test. **E)** Total number of epileptic events during the induction protocol. Ndufs4cKO-vYFP mice n=5, Ndufs4cKO-vr mice n=6. * p<0.05, unpaired t-test. **F)** Total duration of epileptic events. Ndufs4cKO-vYFP mice n=5, Ndufs4cKO-vr mice n=6. ** p<0.01, unpaired t-test**. G)** Maximum severity of epileptic events according to the modified Racine scale. n=6 per group. ** p<0.01, unpaired t-test. **H-K)** Immunofluorescence staining for the microglial marker IBA1 (shown in red) in the GPe from Ndufs4cCT **(H)**, Ndufs4cKO **(I)**, Ndufs4cKO-vYFP **(J)**, and Ndufs4cKO-vr **(K)** mice. Scale bar = 150 µm. **L)** Quantification of IBA1 intensity, expressed in arbitrary units (AU). Ndufs4cCT (n=5), Ndufs4cKO (n=3), Ndufs4cKO-vYFP (n=6), Ndufs4cKO-vr (n=6). * p<0.05, ** p<0.01, indicates significant differences compared to Ndufs4cCT. # p<0.05, indicates significant differences compared to Ndufs4cKO-vYFP. One-way ANOVA test.

To study and compare both the frequency and severity of epileptic events between groups, seizures were induced by increasing the body temperature of Ndufs4cKO-vr and control mice (Oakley et al. 2009; Bolea et al. 2019). Viral vector restoration of NDUFS4 in the GPe of Ndufs4cKO mice (Ndufs4cKO-vr group) led to a significant decrease in epileptic events compared to the control (Ndufs4cKO-vYFP) group (Figure 1D-G). Notably, it was observed that a higher increase in body temperature was required for the rescued mice to develop epilepsy compared to the control group. Moreover, following the complete induction procedure, only 50% of the rescue group animals developed epilepsy, compared to 100% of the control group animals (Figure 1D). Significant differences were found in the total number of epileptic events during the induction protocol. Specifically, during the 10-minute pre-habituation period, 50% of control-injected animals showed epileptic events, while none of the viral rescue animals did. Differences were also observed during the temperature increase period (Figure 1E). Likewise, significant reductions in the total duration and maximum severity of epileptic events were observed between the Ndufs4cKO-vr group compared to the Ndufs4cKO-vYFP group (Figure 1E-G).

Ndufs4cKO mice exhibit overt microglia/macrophage reactivity in the GPe (Chauhan et al. 2022).Therefore, we set to determine whether *Ndufs4* re-expression could alter microglial/macrophage reactivity and infiltration in the in the GPe of cKO mice. The results of immunofluorescence analysis for the microglial protein IBA1 revealed that *Ndufs4* re-expression significantly reduced microglial reactivity in Ndufs4cKO mice compared to uninjected or control-injected animals (Figure 1H-L).

To rule out the possibility that the beneficial effects of the rescue vector, compared to the control vector, were due to serotype-mediated differences in cellular transduction or distinct cellular compartmentalization of the control protein (mitochondrial or cytosolic), we generated a mitochondria-targeted control AAV5 vector by fusing the YFP sequence with the mitochondrial targeting sequence (MTS) for COX8 (AAV-DIO-COX8(MTS)·YFP, Figure S1). Injection of this control viral vector, and the re-expression viral vector, into the GPe of Ndufs4cKO mice (Ndufs4cKO-vmtYFP and Ndufs4cKO-vr mice, respectively; Figure S1I, J) confirmed previous results, showing that *Ndufs4* re-expression in the GPe of Ndufs4cKO mice significantly increased their lifespan (Figure S1K).

Once demonstrated that restoring NDUFS4 in the GPe of Ndufs4cKO mice is crucial for improving their phenotype and reducing epileptic events in this animal model, we set to investigate whether the GPe was not only a necessary and implicated area in the epileptic process in Ndufs4cKO mice, but also sufficient for seizure generation. To that end, we stereotaxically injected viral vectors expressing either functional Cre recombinase (AAV-CRE·GFP) or a non-functional Cre as control (AAV-ΔCRE·GFP) (Bruchas et al. 2011; Fadok, Dickerson, and Palmiter 2009; Quintana et al. 2012) into the GPe of mice carrying two floxed *Ndufs4* alleles (Flox-Ndufs4 mice) (Kruse et al. 2008). Histological analysis of the injections confirmed that the expression of the viral vectors was restricted to the GPe and the reticular thalamic nucleus (RTN; Figure S2A,B). Neither of the Flox-Ndufs4 mouse groups (injected with AAV-CRE·GFP or AAV-ΔCRE·GFP) showed spontaneous seizures or temperature-induced epileptic events, in contrast to Ndufs4cKO animals (Figure S2C). This suggests that *Ndufs4* deletion limited to the GPe (and RTN) is not sufficient to trigger epilepsy or convulsive events. However, while Flox-Ndufs4 mice with specific deletion of *Ndufs4* in the GPe did not exhibit epileptic events, our results demonstrated that this deletion led to marked microgliosis confined to the injection site, a response absent in control mice (Figure S2D-J). This indicates that *Ndufs4* deletion in the GPe induces local microglial reactivity and/or macrophage infiltration.

### *Lhx6*-expressing GABAergic neurons in the GPe show increased vulnerability to mitochondrial dysfunction

To assess the impact of *Ndufs4* deficiency on the abundance and survival of GABAergic neurons in the GPe, we generated a mouse line allowing selective expression of the ribosomal protein RPL22 tagged with hemagglutinin (HA) exclusively in *Gad2*-expressing GABAergic neurons (experimental mice: Ndufs4cKO-HA, and control mice: Ndufs4cCT-HA), enabling the visualization of *Gad2* neurons with the use of an anti-HA antibody in Ndufs4cCT-HA and Ndufs4cKO-HA mice euthanized at PD 60 (Figure 2A, B). A reduction in *Gad2*-expressing neurons was observed in the GPe of Ndufs4cKO-HA mice compared to Ndufs4cCT-HA mice (Figure 2C), particularly in the rostral part (from Bregma -0.22 mm to -0.46 mm, Figure 2D), underscoring the potential localization of a compromised neuronal population in this region.

**Figure 2.**
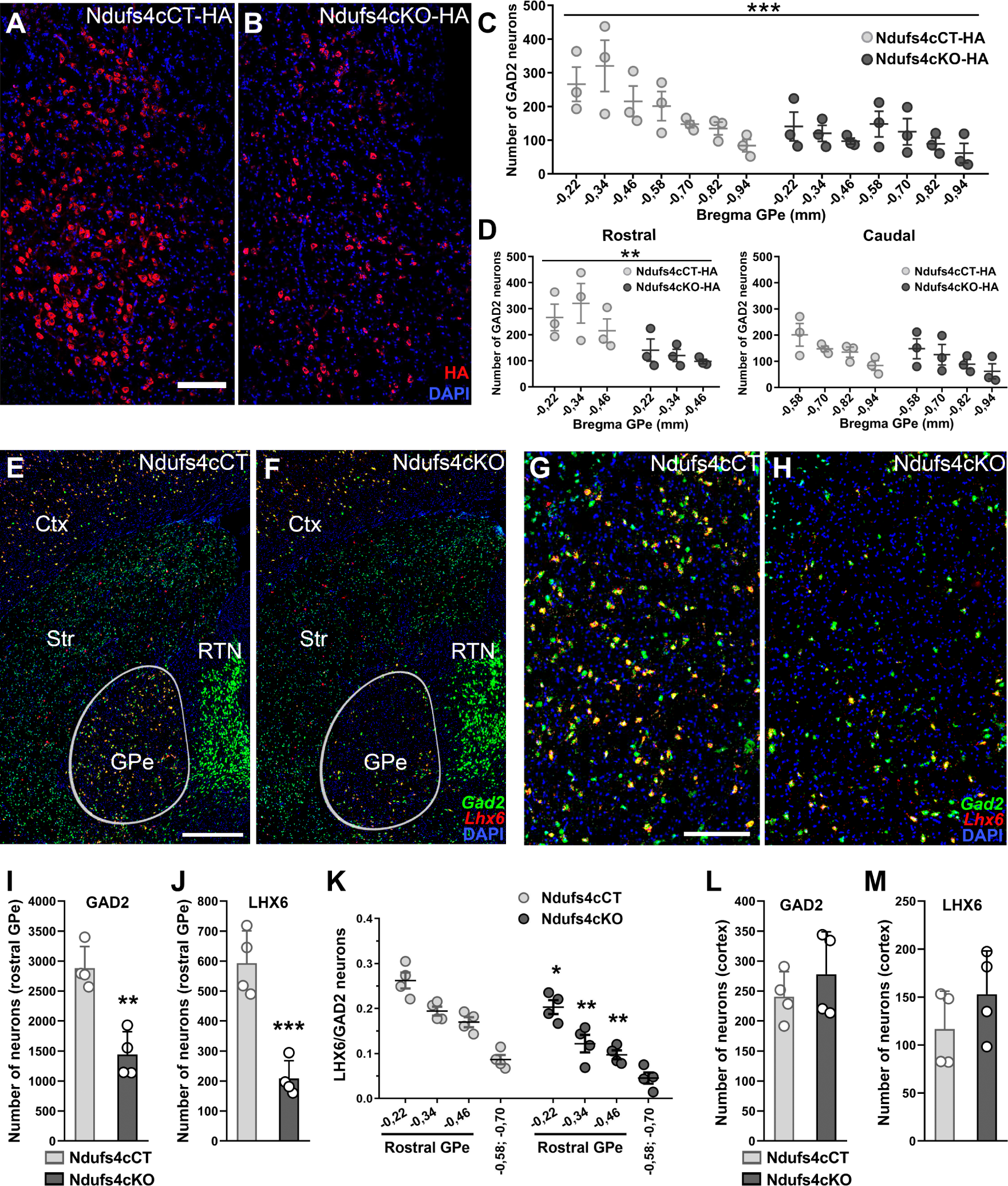
Loss of *Lhx6*-expressing GABAergic neurons in the rostral GPe of Ndufs4cKO mice. **A-B)** Immunofluorescence analysis of the GPe showing HA (red) staining in Ndufs4cCT-HA (A) and Ndufs4cKO-HA (B) mice. Scale bar= 200 µm. **C)** Quantification of HA-positive (GAD2) neurons across the GPe (from Bregma –0.22 mm to –0.94 mm) in Ndufs4cCT-HA and Ndufs4cKO-HA mice. n=3 per group. *** p<0.001, two-way ANOVA. **D)** Quantification of HA-positive (GAD2) neurons in the rostral GPe (from Bregma –0.22 mm to –0.46 mm) and caudal GPe (from Bregma –0.46 mm to – 0.94 mm) from Ndufs4cCT-HA and Ndufs4cKO-HA mice. n=3 per group. ** p<0.01, two-way ANOVA. **E-H**) *In situ* hybridization (ISH) analysis for *Gad2* (green) and *Lhx6* (red) mRNAs in brain sections containing the rostral GPe of Ndufs4cCT (E) and Ndufs4cKO mice (F). Scale bar= 500 µm. Higher magnification images of the rostral GPe of Ndufs4cCT (G) and Ndufs4cKO mice (H). Scale bar=200 µm**. I-J)** Quantification of the total number of *Gad2*-(I) and *Lhx6*-(J) expressing neurons in the rostral GPe (from Bregma –0.22 mm to –0.46 mm) of Ndufs4cCT and Ndufs4cKO mice. n=4 per group. ** p<0.01, *** p<0.001, unpaired t-test. **K)** Proportion of *Lhx6*-expressing neurons relative to the total number of *Gad2*-expressing neurons in the rostral GPe of Ndufs4cCT and Ndufs4cKO mice. n=4 for each group. *** p<0.001, two-way ANOVA. **L-M)** Quantification of the total number of *Gad2*-(L) and *Lhx6*-(M) expressing neurons in the primary somatosensory cortex of Ndufs4cCT and Ndufs4cKO mice (Bregma -0.34 mm) of Ndufs4cCT and Ndufs4cKO mice. n=4 per group.

Approximately 95% of GPe neurons are GABAergic and can be classified into different subtypes based on specific marker expression (Dong et al. 2021), each with distinct projections and anatomical distributions (Abrahao and Lovinger 2018; Dong et al. 2021). Notably, the *Lhx6*-expressing GABAergic subpopulation is more abundant in rostral GPe (Abrahao and Lovinger 2018). *In situ* hybridization (ISH) assays for *Gad2* and *Lhx6* on brain sections of Ndufs4cKO and Ndufs4cCT mice containing the GPe (Figure 2E-H) showed a reduction in both *Gad2*-expressing neurons (Figure 2I) and *Lhx6*-expressing neurons (Figure 2J) in the rostral GPe in Ndufs4cKO mice compared to Ndufs4cCT mice. Moreover, the *Lhx6/Gad2* ratio was also reduced in Ndufs4cKO mice compared to Ndufs4cCT, suggesting increased vulnerability of *Lhx6* neurons in the rostral GPe to *Ndufs4* deficiency (Figure 2K). In contrast, no reduction of *Gad2-* or *Lhx6*-expressing neurons was observed in the cortex of these mice (Figure 2L, M), an area with abundant *Gad2-* and *Lhx6*-expessing neurons, ruling out a global affectation of *Lhx6*-expressing neuronal populations.

### Fatal epileptic events lead to activation of the subthalamic nucleus (STN) in Ndufs4cKO mice

The GPe exhibits predominant projections to various basal ganglia regions, including the striatum, STN, internal globus pallidus (Gpi) and substantia nigra pars reticulata (SNr). In addition, GPe projections also extend to the RTN, cerebral cortex, amygdala, and lateral habenula (Dong et al. 2021). To specifically identify projections from the rostral GPe, stereotaxical injections of a Cre-expressing viral vector (AAV1-CRE-GFP) were performed into the rostral GPe of a tdTomato reporter mice (Ai9 mice) (Madisen et al. 2010). This confirmed pallidal projections to the lateral habenula, STN, and SNr (Figure 3A-C).

**Figure 3.**
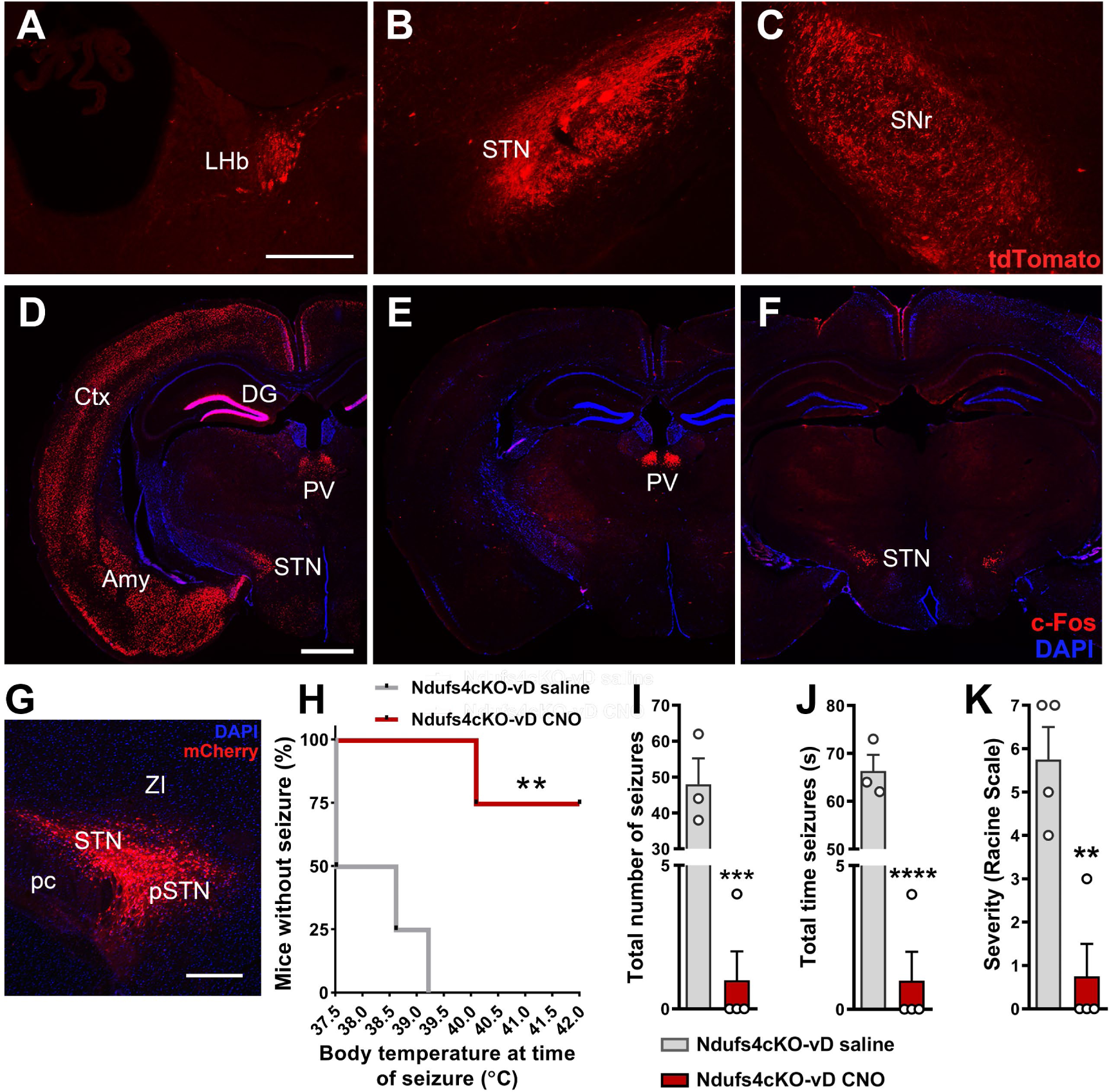
Pallido-subthalamic projections participate in the development of seizures in Ndufs4cKO mice. **(A-C)** TdTomato labeling of brain areas receiving inputs from the GPe: lateral habenula (LHb; A), subthalamic nucleus (STN; B), and substantia nigra pars reticulata (SNr; C). Scale bar= 300 µm. n=3 per group. **(D-F)** Representative images of the brain areas in Ndufs4cKO animals showing c-Fos immunoreactivity following fatal epileptic events. D) Generalized c-Fos expression pattern. E) Restricted c-Fos expression in the paraventricular thalamus (PV). Restricted c-Fos expression in the subthalamic nucleus (STN). DG: Dentate gyrus; Amy: amygdala; Ctx: cortex. Scale bar=1 mm. **(G-K)** Analysis of temperature-induced epilepsy in Ndufs4cKO mice comparing the group with chemogenetic inhibition of the STN (Ndufs4cKO-vD CNO) to the control group (Ndufs4cKO-vD saline). **G)** Representative image of the stereotaxic injection site of the AAV8/2-CaMKIIa-hM4Di·mCherry viral vector in the STN. Scale bar= 400 µm. STN: subthalamic nucleus; pSTN: parasubthalamic nucleus; pc: cerebellar peduncle; ZI: zona incerta. **H)** Percentage of animals without epilepsy relative to body temperature. A value less than 100% at the initial temperature (37.5 °C) indicates that the mouse experienced epileptic events during the habituation period. n=4 per group. ** p<0.01, Log-rank test (Mantel-Cox). **I)** Total number of events during the induction protocol. Ndufs4cKO-vD CNO n=4; Ndufs4cKO-vD saline n=3. *** p<0.001, unpaired t-test. **J)** Duration of epileptic events. Ndufs4cKO-vD CNO n=4; Ndufs4cKO-vD saline n=3. **** p<0.0001, unpaired t-test. **K)** Maximum severity of epileptic events according to the Racine scale. n=4 per group. ** p<0.01, unpaired t-test.

Furthermore, to elucidate the putatively affected and/or involved regions within the epileptogenic circuit of Ndufs4cKO mice, we assessed c-Fos protein expression as a proxy for neuronal activation following epileptic events (Dragunow and Robertson 1987; Morgan et al. 1987). Using this approach, we identified two distinct patterns of neuronal activation in response to fatal epileptic events in Ndufs4cKO animals. Approximately half of the Ndufs4cKO mice exhibited widespread c-Fos staining across various brain regions, including the dentate gyrus of the hippocampal formation, cerebral cortex, STN, and paraventricular thalamus, among others (Figure 3D). In contrast, a second group of Ndufs4cKO animals showed c-Fos staining confined to the paraventricular thalamus (PVT) and STN regions only (Figure 3E-F). Thus, despite the marked difference in overall c-Fos activation, the STN and paraventricular thalamus were the only areas consistently activated across all animals.

### Chemogenetic inhibition of glutamatergic neurons in the STN attenuates temperature-induced epilepsy in Ndufs4cKO mice

Pallidal *Lhx6*-expressing GABAergic neurons represent one of the GPe subpopulations that project to the glutamatergic neuronal populations in the STN (Abdi et al. 2015; Abrahao and Lovinger 2018). Consequently, we hypothesized that inhibiting the STN could mitigate epileptic events in Ndufs4cKO mice.

To achieve chemogenic inhibition of glutamatergic neurons in the STN of Ndufs4cKO mice, an AAV8/2-CaMKIIa-hM4Di·mCherry viral vector was stereotaxically administered into the STN of Ndufs4cKO mice (Figure 3G). Subsequently, to assess the role of the STN in epileptic events following chemogenetic inhibition, mice were subjected to a thermally-induced epileptogenic paradigm (Bolea et al. 2019) on postnatal day 54. After completing the entire induction procedure, only 25% of animals with chemogenetic inhibition (CNO-treated) exhibited epilepsy, in contrast to 100% in the saline group (Figure 3H). In addition, the CNO-treated mice presented a higher threshold for temperature-induced epileptic events, together with a decrease in the frequency, duration, and maximum severity of epileptic events during the induction protocol compared to the saline group (Figure 3I-K).

### Specific loss of *Lhx6-expressing* GABAergic neurons in the GPe is conserved in animals constitutively lacking NDUFS4

Ndufs4cKO mice manifest an overt vulnerability of GABAergic neurons in the rostral GPe to *Ndufs4* deletion, particularly within the *Gad2*/*Lhx6*-expressing subpopulation. Thus, we aimed at validating whether this vulnerability is conserved in animals that constitutively lack NDUFS4 (Ndufs4KO mice), an established model of Leigh syndrome (Kruse et al. 2008; Quintana et al. 2010; Quintana et al. 2012; van de Wal et al. 2022). In agreement with our results in the conditional *Ndufs4* knock-out in GABAergic neurons, *in situ* hybridization assays showed a reduction in the number of *Gad2*-expressing neurons in the rostral GPe in the constitutive knockout group compared to controls (Figure 4A-E). Furthermore, a decrease in the number of *Lhx6*-expressing neurons in the rostral GPe was also observed in the Ndufs4KO mice compared to controls (Figure 4A-D, F). A significant reduction in the *Lhx6*/*Gad2* ratio further underscored a preferential decrease in *Lhx6*-expressing neurons in the rostral GPe in animals lacking *Ndufs4* compared to control mice (Figure 4G). However, no such difference in the number of *Gad2-* or *Lhx6*-expressing neurons was observed in the cortex (Figure 4H-I), an area not predominantly affected in Ndufs4KO mice (Kruse et al. 2008; Quintana et al. 2010; Quintana et al. 2012; van de Wal et al. 2022). These results suggest that the reduction of *Lhx6*-expressing neurons in the rostral GPe is a conserved response to Ndufs4 deficiency, highlighting the pallidal region’s specific vulnerability.

**Figure 4.**
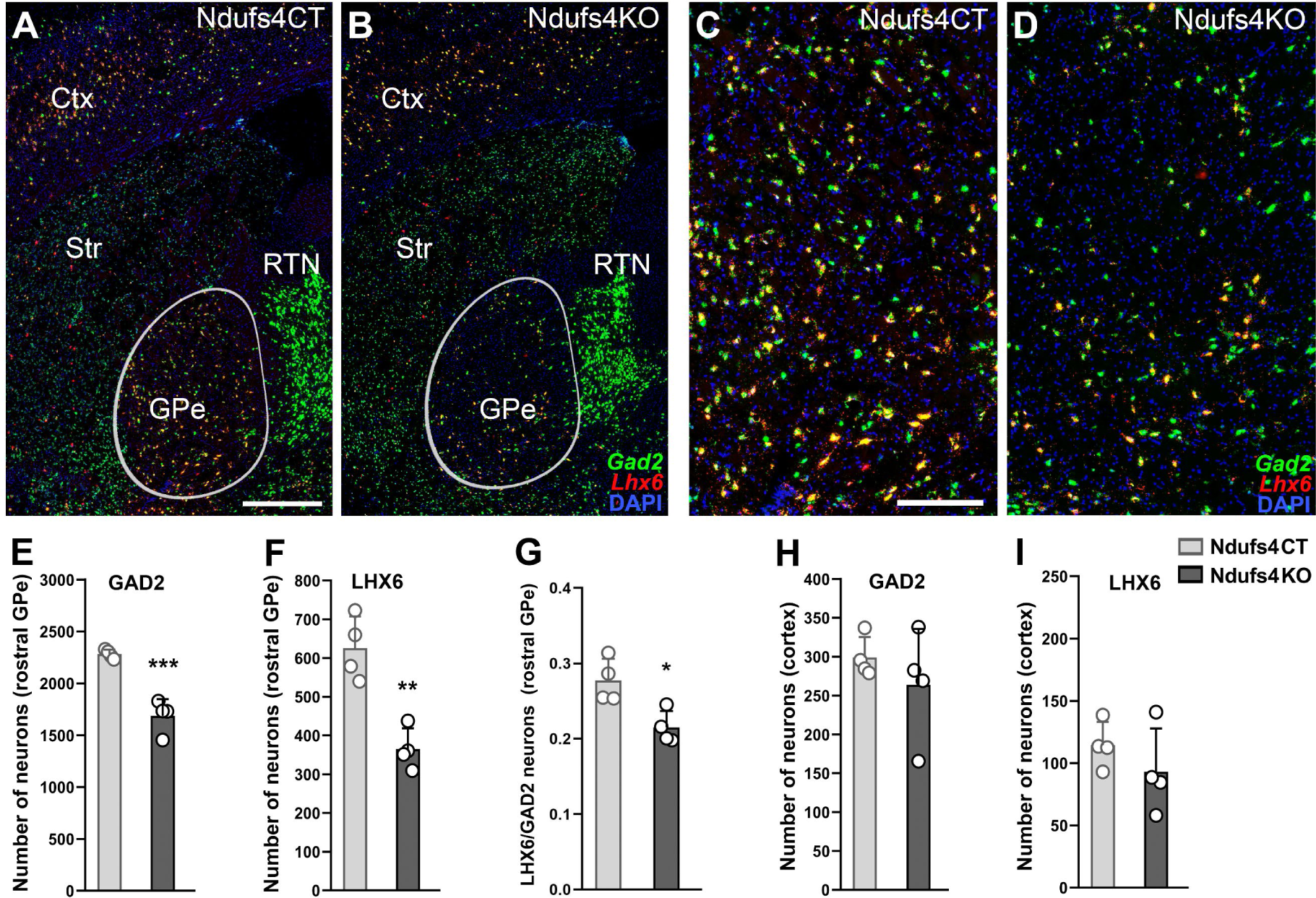
Reduced number of *Gad2-* and *Lhx6-*expressing neurons in the rostral GPe of Ndufs4KO mice. **A-D)** *In situ* hybridization (ISH) analysis for *Gad2* (green) and *Lhx6* (red) mRNAs in brain sections containing the rostral GPe of Ndufs4CT (A, C) and Ndufs4KO (B, D) mice. C-D show magnified views of the GPe in A and B, respectively. **E-F)** Quantification of the total number of *Gad2*-(A) and *Lhx6*-(B) expressing neurons in the rostral GPe of Nfufs4KO and control mice (from Bregma –0.22 mm to –0.46 mm). n=4 per group. ** p<0.01, *** p<0.001, unpaired t-test. **G)** Proportion of *Lhx6*-expressing neurons relative to the total number of *Gad2*-expressing neurons in the rostral GPe of Ndufs4CT and Ndufs4KO mice. n=4 per group. * p<0.05, Two-way ANOVA. **H-I)** Quantification of the total number of *Gad2*-(H) and *Lhx6*-expressing (I) neurons in the primary somatosensory cortex of Ndufs4CT and Ndufs4KO mice (Bregma -0.34 mm). n=4 per group.

### NDUFS4 restoration in external pallidal GABAergic neurons reduces temperature-induced seizures and extends lifespan in Ndufs4KO mice

Ndufs4KO mice exhibit a fatal phenotype associated with reduced lifespan (approximately between postnatal day -PND-50 and PND60) (Quintana et al. 2010; Quintana et al. 2012). Ndufs4KO mice manifest spontaneous epilepsies (Quintana et al. 2010) and are susceptible to thermally induced seizures (Bornstein et al. 2022). Thus, we set to assess the involvement of GPe GABAergic populations in the development of the epileptic phenotype of Ndufs4KO mice. To that end, we first generated a mouse line with a constitutive deletion of *Ndufs4* and concurrent expression of Cre recombinase in GABAergic neurons (Gad2^Cre^,Ndufs4KO mice). These mice subsequently received a stereotaxic injection of a Cre-dependent AAV viral vector expressing either *Ndufs4* or a mitochondria-targeted YFP in the GPe. Re-expression of *Ndufs4* in the GPe increased the survival of the Gad2^Cre^,Ndufs4KO mice (Gad2^Cre^,Ndufs4KO-vr group *versus* Gad2^Cre^, Ndufs4KO-vmtYFP and non-injected Gad2^Cre^,Ndufs4KO mice) (Figure 5A). Analysis of thermal-induced epileptic events during the induction protocols at different disease stages (PND40, 47, and 54) in Gad2^Cre^,Ndufs4KO mice revealed an overall significant decrease in seizure frequency in the Gad2^Cre^,Ndufs4KO-vr group compared to the Gad2^Cre^,Ndufs4KO-vmtYFP and non-injected Gad2^Cre^,Ndufs4KO control groups (Figure 5B). Noteworthy, we observed a differential impact of NDUFS4 restoration during disease progression (Figure 5C-K). In this regard, at PND40, Gad2^Cre^,Ndufs4KO-vr mice presented a higher resistance to thermally-induced seizures, showing approximately a threefold reduction in the number of animals developing seizures compared to Gad2^Cre^,Ndufs4KO-vmtYFP, or Gad2^Cre^,Ndufs4KO mice (Figure 5C). In addition, a reduction in the frequency (Figure 5D), and severity (Figure 5E) of epileptic events was observed in the Gad2^Cre^,Ndufs4KO-vr group. At PND47, Gad2^Cre^,Ndufs4KO-vr mice continued to exhibit resistance to epilepsy induction compared to the Gad2^Cre^,Ndufs4KO-vmtYFP and Gad2^Cre^,Ndufs4KO groups. However, all animals eventually experienced epileptic events (Figure 5F). Likewise, a significant decrease was observed in the total number of epileptic events during the induction protocol in the Gad2^Cre^,Ndufs4KO-vr group compared to both control groups (Figure 5G). However, no differences were noted in the maximum severity of these events (Figure 5H). Finally, at a late stage of the disease, specifically at PND54 (Figure 5I-K), the Gad2^Cre^,Ndufs4KO-vr group continued to show a significant reduction in total number of epileptic events compared to the Gad2^Cre^,Ndufs4KO-vmtYFP and Gad2^Cre^,Ndufs4KO control groups (Figure 5J). Nonetheless, no differences were found in the percentage of mice developing seizures (Figure 5I) or the severity of the events (Figure 5K).

**Figure 5.**
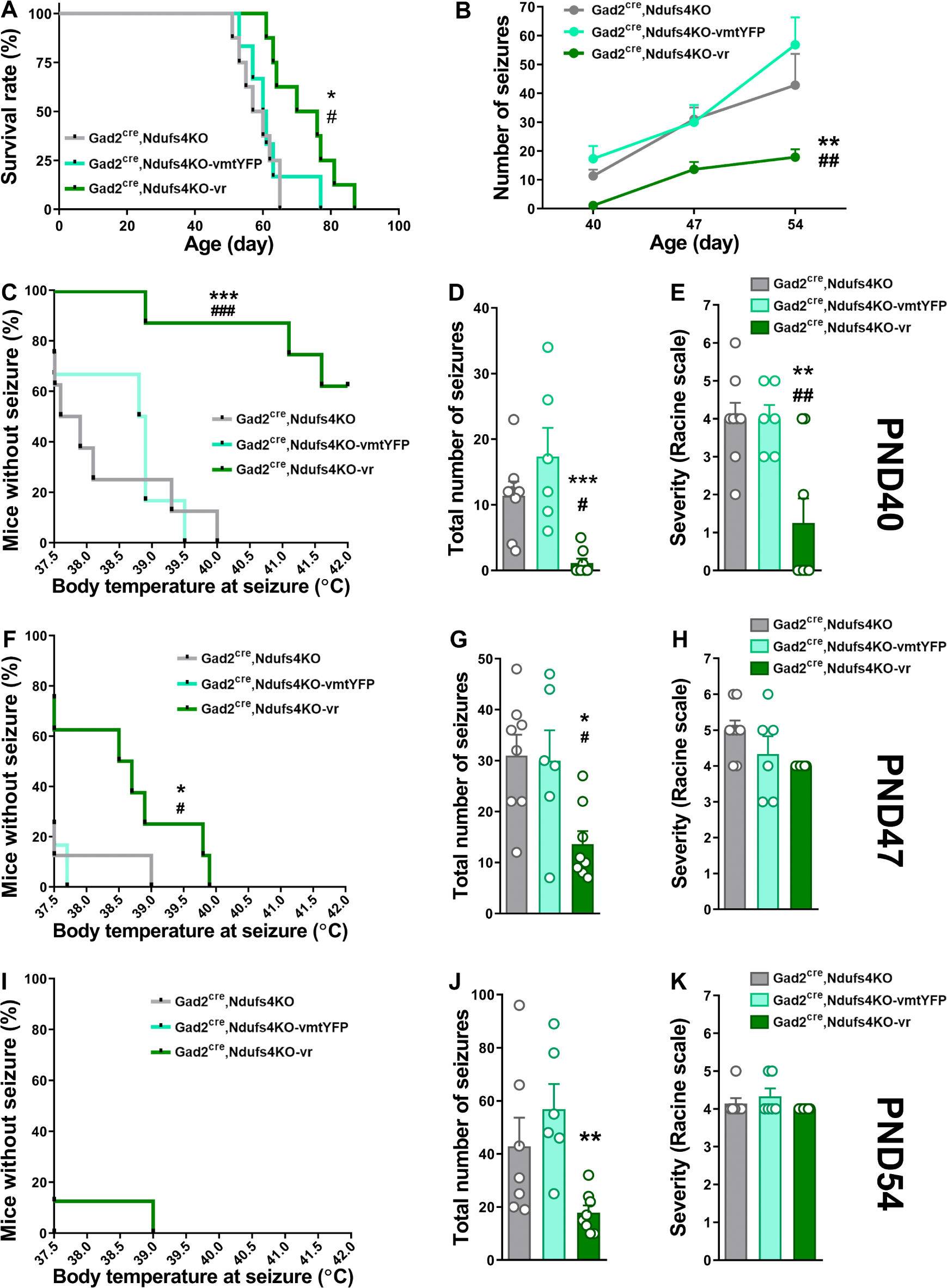
*Ndufs4* re-expression in external-pallidal *Gad2*-expressing neurons extends lifespan and reduces temperature-induced seizures in *Ndufs4*-deficient animals. **A)** Survival curves of Gad2^Cre^, Ndufs4KO (uninjected; n=8), Gad2^Cre^, Ndufs4KO-vmtYFP (control-injected; n=6), and Gad2^Cre^, Ndufs4KO-vr (viral rescue; n=8) mice. # p<0.05 indicates significant differences compared to Gad2^Cre^, Ndufs4KO mice, * p<0.05 indicates significant differences compared to Gad2^Cre^, Ndufs4KO-vmtYFP, Log-rank test (Mantel-Cox). **B)** Temporal progression of total epileptic events during inductions at PND40, PND47, and PND54 in Gad2^Cre^, Ndufs4KO (n=7-8), Gad2^Cre^, Ndufs4KO-vmtYFP (n=6), and Gad2^Cre^, Ndufs4KO-vr (n=8) groups. ## p<0.01 indicates significant differences compared to Gad2^Cre^, Ndufs4KO mice, ** p<0.01 indicates significant differences compared to Gad2^Cre^, Ndufs4KO-vmtYFP mice; Mixed-effects analysis. **C-E)** Effect of NDUFS4 restoration in the GPe on temperature-induced epilepsy in Gad2^Cre^, Ndufs4KO mice at PND40, as assessed by: the percentage of animals remaining seizure-free after increasing body temperature (C), the total number of epileptic events during the induction protocol (D) and the severity of epileptic events according to the Racine scale (E). Gad2^Cre^, Ndufs4KO n=8; Gad2^Cre^, Ndufs4KO-vmtYFP n=6; Gad2^Cre^, Ndufs4KO-vr n=8. ** p<0.01, *** p<0.001, indicates significant differences compared to the Ndufs4KO-vmtYFP group. # p<0.05, ## p<0.01, ### p<0.001, indicates significant differences compared to Gad2^Cre^, Ndufs4KO group. Panel C: Log-rank test (Mantel-Cox). Panels D and E: One-way ANOVA. **F-H)** Effect of NDUFS4 restoration in the GPe on temperature-induced epilepsy in Gad2^Cre^, Ndufs4KO mice at PND47, as assessed by: the percentage of animals remaining seizure-free after increasing body temperature (F), the total number of epileptic events during the induction protocol (G) and the severity of epileptic events according to the Racine scale (H). Gad2^Cre^, Ndufs4KO n=8; Gad2^Cre^, Ndufs4KO-vmtYFP n=6; Gad2^Cre^, Ndufs4KO-vr n=8. * p<0.05, indicates significant differences compared to the Ndufs4KO-vmtYFP group. # p<0.05, indicates significant differences compared to Gad2^Cre^, Ndufs4KO group. Panel F: Log-rank test (Mantel-Cox). Panels G and H: One-way ANOVA. **I-K)** Effect of NDUFS4 restoration in the GPe on temperature-induced epilepsy in Gad2^Cre^, Ndufs4KO mice at PND54, as assessed by: the percentage of animals remaining seizure-free after increasing body temperature (I), the total number of epileptic events during the induction protocol (J), and the severity of epileptic events according to the Racine scale (K). Gad2^Cre^, Ndufs4KO n=7; Ndufs4KO-vmtYFP n=6; Gad2^Cre^, Ndufs4KO-vr n=8. ** p<0.01, indicates significant differences compared to the Gad2^Cre^, Ndufs4KO-vmtYFP group. Panel I: Log-rank test (Mantel-Cox). Panels J and K: One-way ANOVA.

## DISCUSSION

Epilepsy significantly impacts quality of life, with early-onset cases associated with a severe prognosis and reduced life expectancy (El Sabbagh et al. 2010; Desguerre et al. 2014). Epilepsy is a frequent manifestation in MD, a group of severe and untreatable pathologies resulting from mutations in affecting mitochondrial function. However, there are no effective therapies for mitochondrial epilepsies, as they are often resistant to common antiepileptic drugs (Rahman 2019).

The underlying mechanisms of mitochondrial epilepsy remain unclear. Variability in seizure incidence and prevalence across MD has hindered the study of the underpinnings of epilepsy in MD (Desguerre et al. 2014; Rahman 2012; Ticci et al. 2020). In this regard, the generation of animal models of MD developing epilepsy (Olkhova et al. 2023; Nikkanen et al. 2016; Bolea et al. 2019; Bornstein et al. 2022) has paved the way to characterize the mechanisms involved in mitochondrial epilepsy. Among them, studies in a mouse model of Leigh syndrome lacking the complex I subunit NDUFS4, have been instrumental identifying the involvement of GABAergic neurons in the development of epileptic events (Bolea et al. 2019; Manning et al. 2023). However, the circuitry involved in mitochondrial epilepsy remains to be elucidated. Here, we use these validated mouse models to identify the critical role of a pallido-subthalamic circuitry in the mitigation of mitochondrial epilepsy.

Leigh Syndrome is characterized by overt symmetrical lesions in brainstem and basal ganglia (Finsterer 2008), areas particularly vulnerable to *Ndufs4* deficiency in mice (Bolea et al. 2019; Quintana et al. 2012). Among them, electrophysiological alterations in GPe neurons had been shown to precede epileptic events in Ndufs4cKO mice (Bolea et al. 2019). Our results show that *Ndufs4* re-expression in the GPe reduces microglial reactivity and improves the epileptic phenotype observed in both Ndufs4cKO and Ndufs4KO mice. These results highlight that the regulation of epileptic events by the GPe is not restricted to a conditional imbalance of the excitatory/inhibitory inputs, potentially intensifying the epileptic condition (Iizuka et al. 2002). Conversely, selective deletion of *Ndufs4* in the GPe increases local microglial reactivity but is not sufficient to elicit the appearance of epileptic seizures. One possible explanation for the lack of epileptic phenotype is that GPe dysfunction due to *Ndufs4* deletion requires targeting of a larger population of GPe neurons than what is achieved through *Ndufs4* re-expression in rescue experiments. However, this seems unlikely given the broad targeting and the overt microglial/macrophage response observed after *Ndufs4* deletion in the GPe. Thus, our results seem to underscore a role for the GPe in the limitation of the propagation of epileptic events initiated elsewhere. The specific area or anatomical region where epilepsies originate from mitochondrial dysfunction remains unknown. Fos gene product staining studies in Ndufs4cKO mice have revealed neuronal activation in the cortex, dentate gyrus of the hippocampal formation, and the amygdala, among other areas associated with epilepsy in humans and rodents (Cormack et al. 2005; Fournier et al. 2013; Centeno et al. 2014; Menassa, Sloan, and Chance 2017). However, these events may be attributed to synchronization and excitotoxicity associated with epilepsy (Barker-Haliski and White 2015), precluding their direct implication in epilepsy genesis (Manning et al. 2023).

Basal ganglia circuits have been suggested to prevent cortical seizure propagation through feedback mechanisms (Badawy et al. 2013; Vuong and Devergnas 2018; Rektor et al. 2012). Accordingly, Ndufs4cKO mice present increased cortical and hippocampal activity initiated by impaired interneuron function (Manning et al. 2023). Thus, our results suggest that the inhibitory network of basal ganglia is compromised in Ndufs4cKO mice, impairing the control of excitatory neuron activity and leading to the development of epilepsy. In this context, rescuing NDUFS4 in the GPe could potentially restore basal ganglia circuit function, ultimately improving the epileptic phenotype.

Our study has identified, for the first time, the selective loss of *Gad2* neurons in the rostral segment of the GPe in Ndufs4cKO mice, in particular *Lhx6-*expressing neurons, which have been described to be abundant in this area (Abrahao and Lovinger 2018; Dong et al. 2021). Deletion of *Lhx6* has been associated to impaired cortical interneuron migration and differentiation, and seizures (Liodis et al. 2007; Neves et al. 2013; Christodoulou et al. 2022). Thus, it is possible that lack of NDUFS4 may lead to widespread interneuron development. However, no alterations in the numbers of either cortical *Gad2*- or *Lhx6*-expressing interneurons was observed, in line with previous reports (Manning et al. 2023), ruling out a global developmental defect as the cause for the seizures. In this regard, the epileptic phenotype commonly worsens as disease progresses in individuals with Leigh Syndrome (Sofou et al. 2014) and NDUFS4KO mice (Bolea et al. 2019; Bornstein et al. 2022; Quintana et al. 2010). Thus, we hypothesize that progressive degeneration of GABAergic populations in affected areas, such as the GPe, reduces the overall inhibitory tone, with potential implications for epilepsy-associated neurodegeneration in both humans and rodents (Kotloski et al. 2002; Henshall and Murphy 2008).

Our activity-mapping studies highlighted a consistent activation of both the STN and PVT after fatal epileptic events. In mammals, the PVT functions as a stress sensor, activated by physical or psychological stressors (Chastrette, Pfaff, and Gibbs 1991; Cullinan et al. 1995; Spencer, Fox, and Day 2004), suggesting it is a proxy of the stressful response triggered by the epileptic event. In contrast, the STN receives major connections from the GPe (Albin, Young, and Penney 1989; DeLong 1990), being the *Lhx6*-expressing neurons one of its main inputs (Abdi et al. 2015). Supporting this, chemogenetic inhibition of glutamatergic STN neurons is sufficient to reduce the frequency and severity of thermal-induced epileptic events in Ndufs4cKO mice. Collectively, these findings implicate the STN as a critical GPe projection involved in controlling and propagating epilepsies in the context of mitochondrial dysfunction.

Neuromodulation of the STN, such as by deep brain stimulation (DBS), has been shown to significantly alter epileptic activity (Ren et al. 2020) in patients with drug-resistant epilepsy as well as in murine experimental models (Klinger and Mittal 2018; Ren et al. 2020; Wang et al. 2020; King-Stephens 2021; Passamonti et al. 2021; Yan, Ren, and Yu 2022). While DBS can induce activation or inhibition of the targeted area (Miocinovic et al. 2013), our results suggest that inhibiting the STN may be beneficial for controlling epilepsy in patients with MD. Based on our results, we propose that the loss of LHX6 neurons in the GPe leads to reduced inhibitory control in the STN, contributing to the propagation of epilepsy in Ndufs4cKO mice. The STN sends excitatory output to the GPi and SNr, the output nuclei of the basal ganglia (Albin, Young, and Penney 1989; DeLong 1990). Additionally, the STN serves as an input nucleus of the basal ganglia through the hyperdirect pathway, receiving direct glutamatergic connections from motor, premotor, and frontal cortical areas (Nambu, Tokuno, and Takada 2002), and establishing reciprocal connections with various thalamic nuclei and the prefrontal cortex (Lanciego, Luquin, and Obeso 2012). In addition, the STN sends excitatory glutamatergic projections to the GPe, amplifying STN inhibition through a feedback loop (Shink et al. 1996). We hypothesize that the impairment and loss of GPe projection neurons to the STN affects this regulatory mechanism. This leads to overexcitation of the STN, contributing to epilepsy spread, as well as the generation of a state of cerebral synchronicity and excitotoxicity triggering fatal epileptic events (Barker-Haliski and White 2015).

In conclusion, this work underscores a novel role for pallido-subthalamic projections in the development of the epilepsy in the context of mitochondrial dysfunction, suggesting STN inhibition as a potential therapeutic intervention for refractory epilepsy in patients with MD.

## ACKNOWLEDGMENTS

This work was supported by two MINECO Ramon y Cajal fellowships (RYC2019-028501-I; E.S., and RyC-2012-11873; A.Q.), and a pre-doctoral fellowship (PRE2018-083179 to L.S-B.). E.S received funds from MICIU Proyectos I+D+i “Retos Investigacion” (RTI2018-101838-J-I00) and Ministerio de Ciencia e Innovación (MICINN) (PID2019-107633RB-I00 and PID2022-142544OB-I00). A.Q. received funds from the European Research Council (Starting grant 1178 NEUROMITO, ERC-2014-StG-638106), SAF2017-88108-R, 1179 PID2020-114977RB-I00, RTC2019-006825-1, PDC2021-121883-I0, RED2022-134786-T, 38 1180 funded by MICIU/AEI /10.13039/5011000110330 and by MICIU/AEI /10.13039/501100011033 1181 and Next GenerationEU/ PRTR, AGAUR (2017SGR-323, 2021SGR-720), Fundació TV3-La 1182 Marató (202030), and “la Caixa” Foundation (ID 100010434), under the agreement 1183 LCF/PR/HR20/52400018. A.Q is a recipient of an ICREA Academia award.

## METHODS

### Mice

Gad2^Cre^ (*Gad2*-IRES-Cre) mice (Taniguchi et al. 2011) were procured from The Jackson Laboratory (Stock No: 028867). Flox-Ndufs4 mice (Ndufs4^lox/lox^) and Ndufs4^Δ/+^ mice were previously generated by our group (Kruse et al. 2008; Quintana et al. 2010). Both male and female mice of various ages were used in this study, with specific details on the sex and age provided in the figure legends. All mice underwent a minimum of 10 generations of backcrossing and were on a C57BL/6J background.

Mice with a conditional deletion of *Ndusf4* in *Gad2*-expressing GABAergic neurons (Gad2^Cre^, Ndufs4^lox/lox^ or Ndufs4cKO) were obtained by crossing mice with one floxed *Ndufs4* allele and expressing Cre recombinase under the control of the *Gad2* promoter (Gad2^Cre^, Ndufs4^lox/+^) with mice carrying two floxed *Ndufs4* alleles (Ndufs4^lox/lox^). Littermate controls were Gad2^Cre^, Ndufs4^lox/+^ (Ndufs4cCT) mice. In all cases, the genotype of the offspring was determined by PCR analysis using primer sequences previously described (Bolea et al. 2019; Kruse et al. 2008).

Mice were group-housed under a 12:12 h light:dark cycle at 22°C, with *ad libitum* access to rodent chow (Teklad Global Rodent Diet #2014S; Envigo) and water, unless otherwise specified. Sex and age-balanced groups of 2-to 7-month-old mice were used for all experimental procedures, and no sex differences were observed. Following surgeries, animals were individually housed until the completion of all experimental procedures. Sample sizes were determined using power analyses, and the number of animals per group in each experiment (n) is provided in figure legends. All experiments were conducted following the recommendations outlined in the Guide for the Care and Use of Laboratory Animals and were approved by the Animal Care and Use Committee of the Universitat Autònoma de Barcelona and the Generalitat de Catalunya.

### Epilepsy induction

Mouse body temperature was regulated using a rectal temperature probe and a heat lamp connected to a feedback-controlled temperature system (Physitemp Instruments Inc, NJ). In brief, following the method outlined by (Oakley et al. 2009), body temperature was incrementally raised by 0.5°C every 2 minutes until a seizure was induced or a maximum temperature of 42°C was reached. Subsequently, mice were cooled using a small fan.

### Viral Vector Production

Recombinant adeno-associated viral vectors (AAV) were generated in human embryonic kidney (HEK293T) cells with AAV1 coat serotype. Purification involved sucrose and CsCl gradient centrifugations, and the final suspension was prepared in 1x Hanks Balanced Saline Solution (HBSS) at a titer of 2 x 10^9^ viral genomes/μL, following established protocols (Quintana et al. 2012). The AAV preparations were aliquoted and stored at −80°C until stereotaxic injection.

### Stereotaxic Surgery

All surgical procedures were conducted under aseptic conditions. Animal anesthesia was induced and maintained with 5% and 1-1.5% isoflurane/O2, respectively. Analgesia (5 mg/kg ketoprofen; Sanofi-Aventis) and ocular protective gel (Viscotears®, Bausch+Lomb) were administered. Mice were then positioned on a heating pad within a robot-operated, 3-dimensional (stereotaxic) frame (Neurostar) for intracerebral viral vector delivery. Stereotaxic coordinates were normalized using a correction factor (Bregma-Lambda distance/4.21) based on Paxinos and Franklin’s coordinates (Franklin and Paxinos 2013). AAV preparations were bilaterally delivered into the GPe using the following coordinates: antero-posterior (AP), −0.46 mm from Bregma; medio-lateral (ML), ± 2.00 mm; dorso-ventral (DV), −4.00 mm from the skull surface. The injection rate was maintained at 0.1 μL/min (0.5 μL per injection site) using a 5 μL-syringe (Hamilton). AAV preparations were bilaterally delivered into the STN using the following coordinates: antero-posterior (AP), −1.94 mm from Bregma; medio-lateral (ML), ± 1.50 mm; dorso-ventral (DV), −4.50 mm from the skull surface. The injection rate was maintained at 0.1 μL/min (0.5 μL per injection site) using a 5 μL-syringe (Hamilton). After infusion, the needle remained in place for 6 min to allow proper diffusion. Subsequent needle withdrawal was performed at 1 mm/min to minimize off-target viral leakage. Only animals with accurate targeting were included in the experiment.

### Tissue Processing and Immunofluorescence Analysis

Upon CO2 asphyxiation, mouse brains were promptly dissected. The collected brains were fixed overnight in 4% paraformaldehyde (PFA) dissolved in phosphate-buffered saline (PBS) and subsequently cryoprotected using 30% sucrose in PBS. Cryo-sectioning was performed by freezing the brains for 5 minutes in dry ice, followed by sectioning using a freezing microtome. For immunofluorescence, 30-μm free-floating sections for AAV targeting, c-Fos and microgliosis analysis or 20-μm free-floating sections for HA staining and neuron count were treated with a blocking solution containing 10% normal donkey serum (NDS) and 0.2% Triton X-100 in PBS for 1 hour at room temperature. Subsequently, the sections were incubated overnight at 4°C with a primary antibody solution in PBS containing 0.2% Triton X-100 and 1% NDS. The solution included the following antibodies: Anti-GFP (#ab13970, Abcam) at 1:2,000; Anti-HA (#71-5500, Thermo Fisher Scientific) at 1:1,000; Anti-IBA1 (#ab178846, Abcam) at 1:1,000; Anti-TOM20 (#sc-11415, Santa Cruz) at 1:750; Anti-cFOS (#sc-7202, Santa Cruz) at 1:750. After three washes in PBS with 0.2% Triton X-100, the sections were exposed to a secondary antibody solution containing the secondary antibody of interest (1:500; #ab150173, abcam; #A21206, Invitrogen; #A31572, Invitrogen; #A21207, Thermo Fisher Scientific) for 1 hour at room temperature. Following the incubation, sections underwent three 5-minute washes in PBS with 0.2% Triton X-100 and were mounted on slides using DAPI Fluoromount (#17984-24, Electron Microscopy Sciences) before visualization with an EVOS imaging system (Thermo Fisher Scientific) or a Zeiss LSM 700 tracking confocal microscope. Microgliosis was analyzed using the ImageJ software (Fiji v1.52). To determine the number of GABAergic neurons in the GPe, Ndufs4cCT-HA mice (control group) and Ndufs4cKO-HA mice (experimental group) were used. This approach allowed the labeling of GABAergic GAD2 cells in the area of interest using an anti-HA antibody, as described in the previous section. To obtain representative images of the entire GPe, 7 sections of 20 μm thickness were selected at intervals of 100 μm, spanning a total of 0.84 mm within the area of interest (from Bregma -0.22 mm to -1.06 mm). The Zeiss LSM 700 confocal microscope equipped with a 20X objective was used to capture images. A total of 15 images were acquired, with an interval of 1.34 μm between optical sections within the 20 μm thick section. To cover the entire GPe area, 3 to 5 images (depending on the position in Bregma) were taken with a minimum overlap of 10%, and subsequently, a 3D collage was created using the Grid Stitching function and the linear blend fusion method of the ImageJ software (Preibisch, Saalfeld, and Tomancak 2009). These collages contained information on the DAPI signal (corresponding to nuclear labeling) and HA signal (for GAD2 neuron labeling) from the 15 optical sections taken. The counting of GPe cells was performed blindly using IMARIS software (v 9.2, Oxford instruments). The surface mask function with 3D images was used to automate the cell counting based on HA labeling, with the following settings: Maximum cell size: 15 μm (to differentiate closely spaced cells), minimum cell size: 5 μm (to exclude incomplete cell fragments or nonspecific labeling). The mask generated for each cell with these settings had to include DAPI within its boundaries to be counted as valid. The results obtained were manually reviewed to ensure the accuracy and consistency of the automated counting.

### In Situ Hybridization Assays

Mouse brains were snap-frozen in Tissue-Tek O.C.T. compound (Sakura) with dry ice and stored at −80 °C until cryosectioning. Coronal sections (16 µm) containing the rostral GPe were used for RNAscope analysis (Advanced Cell Diagnostics), following the manufacturer’s instructions. The following probes were employed: Mm-Gad2 (#439371, Advanced Cell Diagnostics) and Mm-Lhx6-C2 (#422791-C2, Advanced Cell Diagnostics). All *in situ* hybridization assays were imaged using an OLYMPUS VS200 ASW scanner and analyzed with QuPath open-source software. Cell counting was conducted in GPe rostral sections from Bregma −0,22 mm to −0,46 mm in 3-4 slices/animal, bilateral, (n=3 animals). The number of *Gad2*- and Lhx6-expressing cells was determined using QuPath (Bankhead et al. 2017).

### Statistics

Data are presented as the mean ± SEM. Statistical analyses were performed using GraphPad Prism v9.0 software. Appropriate tests were selected based on the experimental design, as specified in the figure legends. Statistical significance (p<0.05) is indicated in the figure legends. The number of mice (n) refers to the number of animals per group in each experiment. Different cohorts of mice were used for different tests to avoid repeated testing, and no attrition was observed.

