## Supplementary data for "Dysfunctional LHX6 pallido-subthalamic projections mediate epileptic events in a mouse model of Leigh Syndrome"

### Figure S1

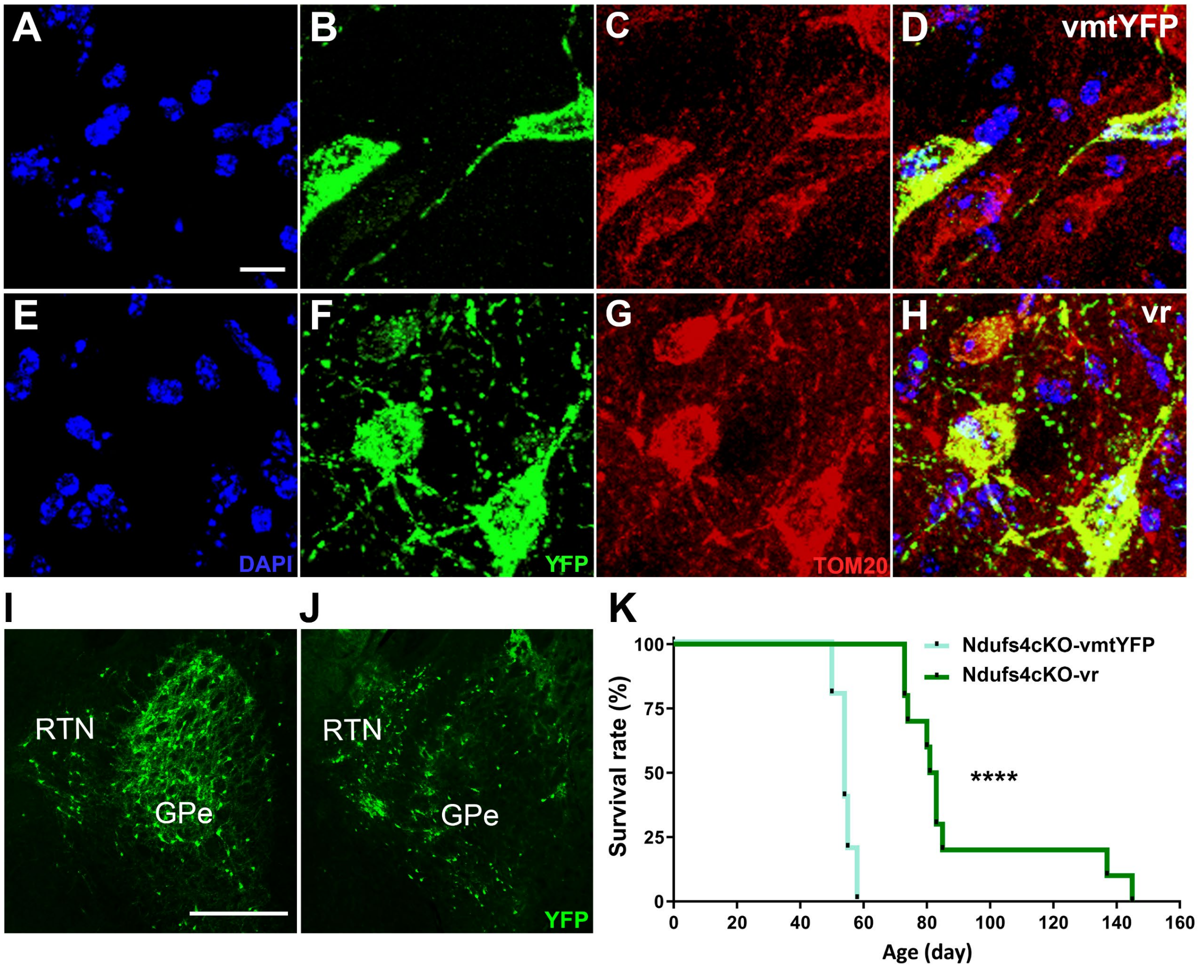


**Figure S1. Validation of the mitochondria-targeted control viral vector and confirmation of lifespan extension following *Ndufs4* re-expression in the GPe of Ndufs4cKO mice. A-H)** Immunofluorescence analysis of Ndufs4cKO mice following stereotactic injection into the GPe with either the AAV5-DIO-COX8(MTS)·YFP viral vector (Ndufs4cKO-vmtYFP mice; A-D) or the AAV5-DIO-NDUFS4·YFP viral vector (Ndufs4cKO-vr mice; E-H). Panels show: nuclei stained with DAPI (A,E), YFP (B, F) and TOM20 (C, G) staining. An overlay of all stainings shows mitochondrial localization of vmtYFP (D) and NDUFS4·YFP (H). Scale bar = 10 μm. **I-K)** *Ndufs4* re-expression in the GPe extends the lifespan of Ndufs4cKO Mice**.** Images show stereotactic injection into the GPe of the rescue AAV5-DIO-NDUFS4·YFP (I) or control AAV5-DIO-COX8(MTS)·YFP (J) viral vectors. Graph (K) shows lifespan extension in Ndufs4cKO-vr mice (n=10) compared to Ndufs4cKO-vmtYFP mice (n=5). RTN: Reticular thalamic nucleus, GPe: Globus pallidus externus. Scale bar = 400 μm. $p<0.0001$, Log-rank test (Mantel-Cox).

### Figure S2

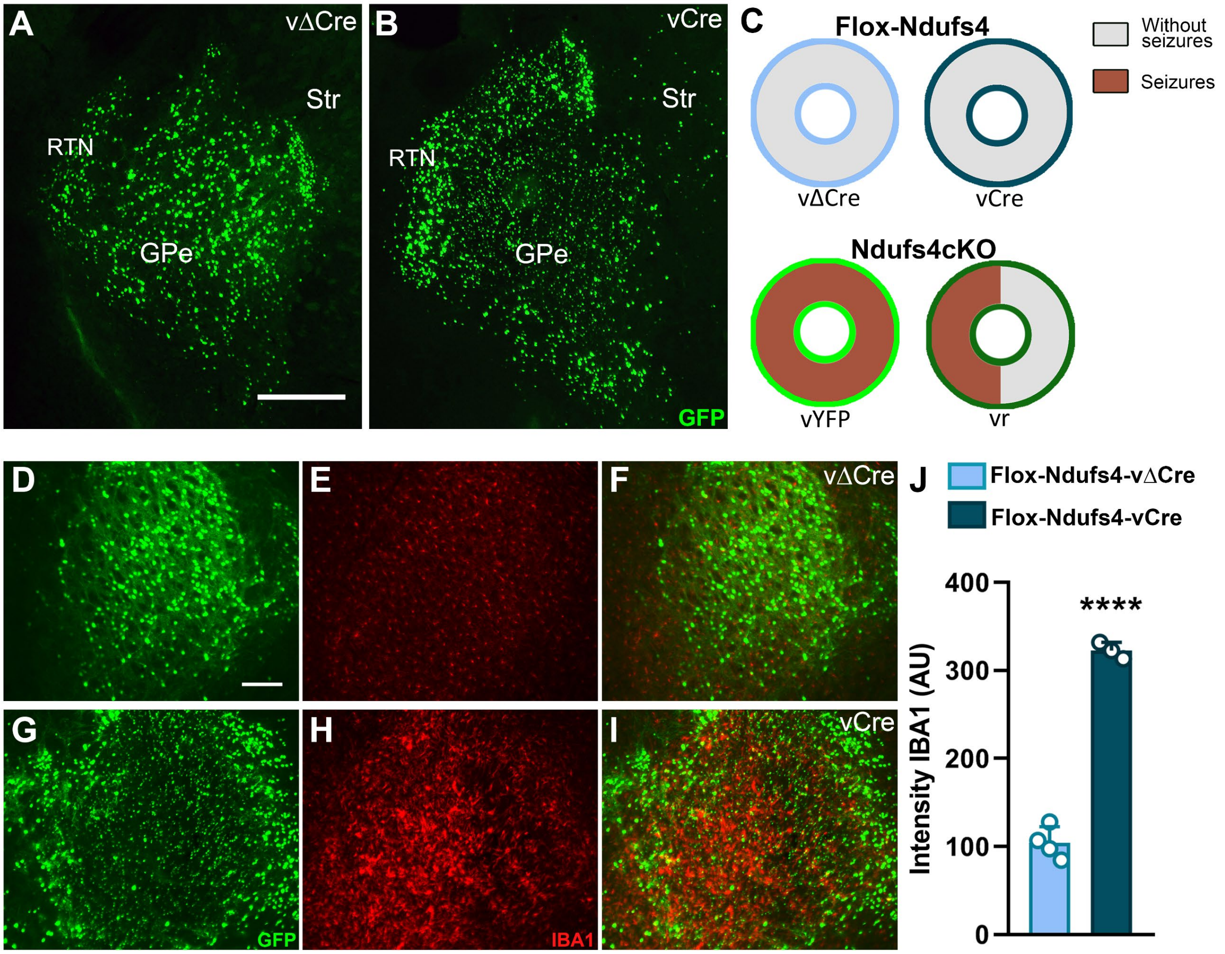


**Figure S2. Effect of acute *Ndufs4* deletion in the GPe on seizure incidence and microglial reactivity. A-B)** Representative images showing transduction efficiency in the GPe of the AAV1-ΔCRE·GFP (vΔCre; A) and the AAV1-CRE·GFP (vCre; B) viral vectors. **C)** Percentage of epileptic events during the induction protocol in Flox-Ndufs4 and Ndufs4cKO mice. Flox-Ndufs4 mice with vΔCre or vCre n=8; Ndufs4cKO mice with vΔCre or vCre n=6; Scale bar = 400 µm. RTN: Reticular thalamic nucleus, GPe: Globus pallidus externus, Str: striatum. **D-J)** Immunofluorescence analysis for IBA-1 (red) and GFP (green) showing microgliosis following Cre-mediated *Ndufs4* deletion in neurons of the GPe of Flox-Ndufs4 mice. Panels D and G show localization of the GFP fluorescent protein from the AAV1-ΔCRE·GFP or the AAV1-CRE·GFP viral vectors, respectively. Panels E and H show expression of the microglial marker IBA1 in the GPe of Flox-Ndufs4 mice transduced with AAV1-ΔCRE·GFP or AAV1-CRE·GFP, respectively. Panels F and I show the overlay of both stainings. Graph in J shows a quantification of IBA1 intensity in Flox-Ndufs4 mice injected with AAV1-ΔCRE·GFP (Flox-Ndufs4-vΔCre; n=4) or AAV1-CRE·GFP (Flox-Ndufs4-vCre; n=3). **** $p<0.0001$, unpaired t-test.
